# Hydrogen Cyanamide Causes Reversible G2/M Cell Cycle Arrest Accompanied by Oxidation of the Nucleus and Cytosol in *Arabidopsis thaliana* Root Apical Meristem Cells

**DOI:** 10.1101/2022.10.20.512991

**Authors:** Yazhini Velappan, Ambra De Simone, Santiago Signorelli, John A Considine, Christine H Foyer, Michael J Considine

## Abstract

Hydrogen cyanamide (HC) is known to stimulate the production of reactive oxygen species (ROS) and also alter growth through modification of the cell cycle. However, the mechanisms by which HC alters cell proliferation and redox homeostasis are largely unknown. This study used roGFP2 expressing Arabidopsis seedlings to measure the oxidation states of the nuclei and cytosol in response to HC treatment. The Cytrap dual cell cycle phase marker system and flow cytometry were used to study associated changes in cell proliferation. HC (1.5mM) reversibly inhibited root growth during a 24h treatment. Higher concentrations were not reversible. HC did not synchronize the cell cycle. In contrast to hydroxyurea, HC caused a gradual accumulation of cells in the G2/M phase and decline of G1/S phase cells 16 to 24h post-treatment. This was accompanied by increased oxidation of both the nuclei and cytosol. Taken together, HC impairs proliferation of embryonic root meristem cells in a reversible manner through restriction of G2/M transition accompanied by increased oxidative poise.

## Introduction

The cellular reduction oxidation (redox) potential regulates cell biochemistry and hence plays a key role in tuning plant development to the local environmental conditions (Sánchez-Fernández *et al*., 1997; Foyer and Noctor, 2011; Boguszewska-Mańkowska *et al*., 2015). A network of interactions between reactive oxygen species (ROS), antioxidants, phytohormones and regulatory proteins act coordinately to control of plant growth (Foyer and Noctor, 2005; 2011). In particular, changes in cellular redox status in response to external cues plays an important role in regulating cell division in the root and shoot meristems (Diaz-Vivancos *et al*., 2010; De Simone *et al*., 2017). Inter-compartmental transport and sequestration of glutathione influences the transition of cells through key cell cycle checkpoints, G1 and G2 by regulating the cellular redox state (Diaz-Vivancos *et al*., 2010; De Simone *et al*., 2017). The nucleus and cytoplasm have similar levels of GSH before entry into the cell cycle and this equilibrium is restored again during G2-M phase of the cell cycle with the redox state being highly regulated during the G1 and G2 checkpoints (Diaz-Vivancos *et al*., 2010; De Simone *et al*., 2017). Roots are particularly vulnerable to direct interaction with phytotoxins and as such are excellent models of plant developmental plasticity in response to stress. Hence the alteration of growth in roots in response to different oxidants and phytochemicals has been widely studied (Zhang *et al*. 2009; Cools *et al*., 2010; Ding *et al*., 2010; Soltys *et al*., 2011, 2012; Tsukagoshi et al., 2010).

Hydrogen cyanamide (HC) is widely used in the horticulture industry to trigger the resumption of growth following dormancy. It is known to repress mitochondrial aldehyde dehydrogenases (Maninang et al., 2015), alter energy metabolism and gene transcription, and trigger an oxidative burst of ROS, subsequently leading to changes in redox homeostasis allied to glutathione state changes (GSH to GSSG) which may in turn regulate cell division (Soltys *et al*., 2011; 2012; Vergara *et al*., 2012). However, whether and how HC alters proliferation of cells via associated alteration in cellular redox state is unknown. A recent study involving hydroxyurea (HU)-treated embryonic roots of Arabidopsis established that progression of cells through the cell cycle is controlled by alterations in cellular redox homeostasis. The redox potential of both the nucleus and cytosol was determined using the redox-sensitive GFP (roGFP) reporter in Arabidopsis seeds (De Simone *et al*., 2017), and showed that depletion of the soluble antioxidant ascorbate disrupted the intracellular redox flux and rhythm of the cell cycle.

The commonality in the ROS mediated regulation of proliferation in root and shoot apical meristems (Tsukagoshi *et al*., 2010; Zeng *et al*., 2017) and the complex nature of the shoot apical meristem inspired the use of roots as a model system to add to the existing knowledge of the mode of action of HC. Hence this study explored the effect of HC on physiological growth and cellular redox homeostasis in relation to alterations in cell proliferation in embryonic roots of *Arabidopsis thaliana*. The insight obtained from this study may later be transferred to other meristematic systems.

## Results

### Hydrogen cyanamide causes reversible effect on embryonic root growth of Arabidopsis thaliana

HC is phytotoxic at high concentrations (Shulman *et al*., 1986; Soltys *et al*., 2011; 2012). At low concentrations, however, HC relieves dormancy and promotes growth in perennial buds (Or et al., 1999; Shulman *et al*., 1986). Although several studies have implicated the involvement of redox regulation in dormancy release by HC, the regulation of growth by HC has not been studied at the cellular level. Hence this study was designed to examine the influence of HC on cell proliferation and redox state. A dose-response experiment was first carried out to determine the concentration of HC that caused reversible perturbation in root growth on short-term treatment without having any phytotoxic effect, analogous to 3mM HU concentration used in an earlier study to determine the effect of cellular redox regulation on status of cell cycle in embryonic root meristems (De Simone *et al*; 2017).

The concentration of HC was optimized through observation of the effect of various concentrations of HC on rate of root growth at the end of 24h treatment and recovery of root growth after 24h of treatment in comparison to 3mM HU and untreated control (Figure 5). 2d old seedlings grown in the dark at 21°C were treated with a range of concentrations of HC (1.5, 2 and 5mM) or 3mM HU for 24h under the same conditions and later released from the treatment by transferring them to HC or HU free media and grown for a further 48h at 21°C in the absence of light. The corresponding control seedlings were also grown under similar conditions. Root length measurements were made immediately after release from treatments (0h), 24h after release (24h) and 48h post-release from treatment (48h) (Figure 5).

Treatment with 1.5mM HC and 3mM HU for 24h caused a reduction in root growth compared to the control. After 24h of treatment with 1.5mM HC and 3mM HU, roots had a growth rate of 45% and 72% of the control roots respectively with the decrease in growth rate by HU not being statistically significant compared to the control (Figure 5). At these concentrations, HU treated seedlings had a relatively higher root growth rate similar to the control in comparison to HC treated seedlings. After 24h of recovery from these treatments, both HC and HU treated roots recovered >50% of the control root growth rate; although the difference between the treated and control roots was not statistically significant 48h after recovery, HC treated roots exhibited recovery comparable to the control reaching 96% of the control root growth rate (Figure 5). However, HU treated roots only recovered 65% of the control root growth. This indicates that short-term treatment with HC at 1.5mM causes a reversible restriction of root growth fairly similar to HU. Moreover, based on the rate of recovery, it has negligible phytotoxicity when treated for 24h, which is the time period of treatment used in this study (Figure 5). 2mM HC had analogous effect on root growth as 1.5mM HC at the end of 24h treatment. Conversely, it only recovered 36% of the control root growth rate 24h after release from the treatment, similar to 5mM HC (Figure 5). Hence 2-5mM HC was considered phytotoxic to root growth under the treatment conditions used in this study. Lower concentrations of HC were not tested, as previous studies on tomato roots have shown that roots showed perturbations in cell division only after 3d of treatment with 1.2mM HC (Soltys *et al*., 2012). Hence, taking into account the smaller size of Arabidopsis roots in comparison to tomato roots, this study started the optimization with a slightly higher concentration of HC (1.5mM) for a shorter duration of 24h to observe pronounced changes in cell cycle without having any phytotoxic effect on growth.

### Hydrogen cyanamide decreases root growth of Arabidopsis thaliana seedlings in a time-dependent manner

Following the optimization of HC concentration and studying the dose-dependent effect of HC, the time-dependent effect of 1.5mM HC on root growth in comparison to the control was studied. 2d old seedlings germinated in the dark at 21°C were transferred to ½MS media supplemented with and without 1.5mM HC and grown for 48h under the same growth conditions. Root growth was measured immediately after the transfer (0h) (Figure 6), 24h and 48h after the transfer (24h and 48h respectively) (Figure 6). Roots of control (untreated) seedlings grew well during the whole treatment period (Fig 7A) maintaining a uniform growth rate of 1.4 and 1.7mm/d by 24 and 48h after treatment respectively (Figure 6B). However, seedlings treated with HC grew shorter roots than the control with a progressive 0.5 and 1.4mm/d decline in root growth rate in comparison to the control after 24 and 48h of HC treatment (Figure 6B). HC caused a 67% reduction in root growth rate after 48h treatment compared to after 24h treatment causing greater reduction of root growth rate with increase in treatment duration. However, surprisingly, the HC treated seedlings developed shorter roots compared to the control throughout the study as can be seen in Figure 6A. On the whole, HC treated seedlings produced shorter roots compared to the control with its effect being more pronounced with longer treatment time.

### Hydrogen cyanamide treatment of embryonic root tips causes gradual accumulation of G2/M phase cells

Cell cycle progression was monitored both *in-vitro* and *in-vivo* in the absence and presence of HC in Arabidopsis embryonic root tips using flow cytometry and Cytrap marker system. Untreated control root tips maintained around 67% G1, 8% S and 25% G2 cells during 24h of treatment showing no significant change in the proportion of cells in each phase of the cell cycle (Figure 7). Untreated root tips of Cytrap seedlings showed undetectable fluorescence when excited at 488nm (G2/M phase) and 559nm (S/G2 phase) during the 24h of treatment (Figure 8, Supplementary Figure S2) indicating the asynchronous nature of cell cycle progression in the proliferation zone of control root tips. However, there was significant difference in cell cycle status of HC treated root tip cells at 16 to 24h of treatment. The proportion of cells in G1 phase showed a significant decrease from >65% during the earlier time points to 55% at 24h. There was around 10% decrease in G1 cells compared to the control 24h post-treatment with HC. This decrease in G1 cells was accompanied by a considerable increase in G2 phase cells at 24h (8%) of HC treatment (Figure 7). Similarly, HC treated root tips of Cytrap seeds showed a gradual increase in fluorescence when excited at 488nm (G2/M phase) from 16h to 24h of HC treatment (Figure 8). These results indicate an accumulation of G2/M phase cells after 16h HC treatment, with corresponding decline in G1/S phase cells (Figure 7, 8). However, no change was detected in the S/G2 (magenta coloured) channel during the 24h of HC treatment compared to the control (Figure 8) and in the distribution of S phase cells (Figure 7). Taken together, these data suggest that treatment of Arabidopsis embryonic root tips with 1.5mM HC for 24h prolongs cell cycle at G2/M phase. However, this data requires further validation through analysis of expression pattern of cell cycle related genes at various time points after treatment with HC in comparison to untreated control.

### Hydrogen cyanamide triggers a higher cellular oxidation compared to hydroxyurea

Fluorescence ratios determined in the nuclei and cytosol of embryonic root proliferation zone cells of germinating Arabidopsis roGFP2 seeds treated with 1.5mM HC and 3mM HU and untreated control was used to calculate the degree of oxidation, which in turn was used to calculate glutathione redox potentials. Untreated control and HU treated root proliferation zone cells had similar mean glutathione redox potentials of −296.8±0.9 and −297.4±1.1mV in the nuclei and −294.1±1.4 and −295.2±1.6mV in the cytosol respectively over 24h treatment (Table 1). However, the nuclei and cytosol of HU treated cells were relatively less oxidized (21% in the nuclei and 24% in the cytosol) compared to the control cells (23% in the nuclei and 26% in the cytosol) (Table 1). The HC treated cells were most oxidized during the 24h period of treatment with glutathione redox potentials of −291.4±1.9 and −289.6±1.4mV and oxidation degree of 31% and 32% in the nuclei and cytosolic compartments respectively (Table 1). On an average, HC treated cells were 5-10% more oxidized than the HU treated and control cells (Table 1). Moreover, cytosol was rather more oxidized than the nuclei in the embryonic root proliferation zone cells irrespective of the treatment (Table 1).

**Table 1.** Glutathione based average cellular redox potential and oxidation degree for control (untreated), HC and HU treated proliferation zone cells of *Arabidopsis thaliana* embryonic roots over 24h treatment period. (Data represent mean values ± SEM; n ≥ 3)

| Cellular compartment | Average cellular redox potential (mV) |  |  | Average degree of oxidation (%) |  |  |
| --- | --- | --- | --- | --- | --- | --- |
|  | *Control* | *HC* | *HU* | *Control* | *HC* | *HU* |
| *Nuclei* | -296.8 ± 0.9 | -291.4 ± 1.9 | -297.4 ± 1.1 | 22.90 ± 2.3 | 31.22 ± 1.6 | 21.35 ± 1.3 |
| *Cytosol* | -294.1 ± 1.4 | -289.6 ± 1.4 | -295.2 ± 1.6 | 26.31 ± 1.6 | 32.48 ± 1.4 | 24.44 ± 1.8 |

### Accumulation of G2/M cells is accompanied by increased oxidation in hydrogen cyanamide treated root tips

Treatment with HC triggered an oxidation event in the cytosol immediately after treatment (2h) increasing the redox potential to −285.74±1.1mV and oxidation degree to 37% respectively, in comparison to the untreated control with a much lower redox potential of < −300mV and oxidation degree of nearly <20% in both nuclei and cytosol (Figure 9A,B; Figure 10A,B). HU treatment caused similar increase in redox potential and oxidation degree in the cytosol (−288.53±0.7mV and 32% respectively) 2h after treatment but only after 4 and 8h of treatment in the nuclei (ca. −289mV and 32% respectively) (Figure 9C,D; De Simone *et al*., 2017). There was no significant change in redox potential or oxidation degree of the nuclei until 12h post-treatment with HC (Figure 9A,B; Figure 11A,B). During the first 8h of treatment, most of the cells in the proliferation zone of HU synchronized root tips were considered to be in G1/S phase (Figure 9C,D; De Simone *et al*., 2017; Cools *et al*., 2010). No such synchronization of cell cycle was observed in HC treated root tip cells (Figure 8).

Redox potential of HC treated cells varied from −303 to −282mV in the nuclei and −298 to −281mV in the cytosol (Figure 9A,B). The HU treated cells remained comparatively less oxidized with nuclei and cytosolic redox potentials ranging from −303.60 to −289.64mV and 304.997 to 286.64mV respectively (Figure 9C,D; Figure 10C,D). The nuclei and cytosol of the control was relatively more reduced than that of HC but not HU treated cells (Table 1). The redox state of nuclei and cytosol of HC treated cells did not differ significantly in comparison to the control in the first 12h of treatment (Figure 9A,B; Figure 10A,B). However, after 12h of treatment with HC, the nuclei and cytosol became highly oxidized, as opposed to the significantly reduced untreated control cells during this period (Figure 9A,B; Figure 10A,B). The degree of oxidation in the nuclei and cytosol remained greater than the control in HC treated cells as opposed to HU treated cells that remained more reduced than the control in the final hours of treatment (12-24h; Figure 10) when the HU synchronized cells were considered to be in G2 and M phases of the cell cycle (De Simone *et al*., 2017; Cools *et al*., 2010) and HC treated cells showed a progressive increase in G2/M cells (Figure 7, 8). Overall, a higher degree of oxidation was observed in both the cellular compartments studied, during early hours of HU treatment when G1 and S phase cells were predominant. Conversely, a lower degree of oxidation was observed during the final hours of HU treatment when majority of cells accumulated in G2 and M phases of the cell cycle (De Simone *et al*., 2017). HC caused gradual accumulation of G2/M cells in the later hours of the 24h HC treatment (Figure 8). Contrary to the redox regulation by HU, HC stimulated higher degree of oxidation in the final hours of treatment parallel to the accumulation of G2/M cells, in addition to an early oxidation event 2h after HC treatment (Figure 7, 8, 10)

## Discussion

HC is an allelochemical commonly used in agriculture to relieve bud dormancy and promote growth (Shulman *et al*., 1983). It is known to act through the generation of ROS, mainly H_2_O_2_, leading to alterations in redox homeostasis associated with the oxidation of glutathione (GSH to GSSG), which may regulate cell proliferation (Vergara *et al*., 2012). However, if and how HC alters cell proliferation through associated changes in cellular redox state is unknown. The complexity of the meristems in other organs makes it difficult to study the mode of action of HC at the cellular level. Hence, this study used Arabidopsis embryonic root system to study the effect of HC on cell cycle and redox at the cellular level.

Regardless of its widespread use as an agrochemical, the effectiveness of HC largely depends on the concentration of application with higher doses being detrimental to growth (Fuchigami and Nee, 1987; Siller-Cepeda *et al*., 1992; Soltys *et al*., 2011; 2012). In this study, 1 day treatment of Arabidopsis seedlings with 1.5mM HC caused ca. 40% reduction in root growth rate compared to the untreated control, consistent with the changes observed in roots of tomato treated with 1.2 mM HC (Soltys *et al*., 2012), onion treated with 2mM HC (Soltys *et al*., 2011), maize treated with 3mM HC (Soltys *et al*., 2014) and lettuce treated with 10μM HC (Kamo *et al*., 2003). Moreover, continuous treatment of Arabidopsis seedlings with the same concentration of HC for 2 days caused 84% decline in root growth rate compared to the untreated control in this study. Interestingly, tomato roots treated with 1.2mM HC only showed 50% reduction in root growth rate compared to the control after 3 days of treatment (Soltys *et al*., 2012), analogous to 3mM HC treated maize roots. This difference in sensitivity to HC could be attributed to the relatively thinner root system of Arabidopsis, thus increased surface area to volume ratio. Prolonged treatment (48h) with 1.5mM HC caused a decrease in root growth compared to the short-term treatment (24h) similar to observations made in tomato in which shrinkage of only root tips but not the distal root segments was observed after 3 days of HC treatment (Soltys *et al*., 2012). The authors ascribed this to earlier cellular differentiation following the end of mitosis rather than to variations in cell length. The same could be true for Arabidopsis roots in this study. However, this needs further validation. The effects of short-term HC treatment on root growth were completely reversible at 1.5mM concentration as opposed to higher concentrations used in this study in agreement with previous observations in tomato roots treated with 1.2mM HC (Soltys *et al*., 2012). However, Arabidopsis roots recovered to control levels 2 days after release from treatment as opposed to tomato roots which required 5 days to reach control levels (Soltys *et al*., 2012). This could perhaps be due to the difference in growth medium, physiological state of the seedlings or their ability to recover from stress. Onion roots treated with similar concentration of HC (2mM) for short-term showed increased growth after recovery from treatment indicating the growth promoting effect of HC at low dosage. Similar growth promoting effects were also observed in lettuce roots treated with low concentrations of rabdosin B (Ding *et al*., 2010). However, such induction of growth was not observed in this study at the lowest concentration used (1.5mM). This could be because this concentration was not low enough to have a growth promoting effect. Therefore, the inhibitory effect of HC in Arabidopsis root growth is dose and time dependent with its effect being more pronounced and irreversible at higher doses and/or when treated for longer durations analogous to earlier observations (Soltys *et al*., 2012; 2014).

Arabidopsis seedlings treated with 3mM HU for short-term did not show significant reduction in root growth rate compared to the control, in agreement with earlier studies by De Simone (2016). HU had a slightly higher root growth rate compared to 1.5mM HC treated seedlings during short-term treatment. However, 2 days after release from treatments, only roots treated with 1.5mM HC were able to fully recover their growth to untreated control levels. The previous study by De Simone (2016) did not observe the recovery effect of 3mM HU, however no phytotoxicity was evident based on staining for cell viability. Hence, this delay in recovering control levels of growth even in the absence of phytotoxicity needs to be explored further.

Earlier studies report that HC mediated reduction in root growth is caused by perturbations in division of cells at the meristematic region (Soltys *et al*., 2012). In this study, treatment with 1.5mM HC for short-term (24h) caused significant alterations in cell cycle status only from 16h of treatment as opposed to HU which alters cell cycle status immediately after treatment causing an accumulation of S phase cells (Cools *et al*., 2010; De Simone, 2016). HC treatment caused a gradual build-up in population of G2 phase cells with an accompanying decline in G1 phase cells implying a gradual decline in dividing cells analogous to observations made in tomato roots treated with 1.2mM HC for 3 days (Soltys *et al*., 2012). This is in contrast to the observation in onion roots treated with 2mM HC, which did not show any changes in the distribution of cells in various cell cycle phases after short-term treatment (Soltys *et al*., 2011). This result was further backed up by observations using the Cytrap marker system which also suggest an accumulation of G2/M phase cells from 16h till the end of treatment. However, the decline in G1 population observed using flow cytometry, could not be verified by this marker system. Moreover, no significant change in S/G2 phase was detected during the 24h HC treatment. This is in contrast to HU treated roots which showed a clear increase in S/G2 cells 5-10h after treatment in an earlier study (De Simone, 2016). This effect of HU is due to synchronization of the cell cycle by a transient G1/S arrest analogous to results reported by Cools *et al*. (2010). The replication restriction imposed by HU in the earlier study was overcome within the first 5-6h of treatment commencing the first cycle of DNA replication following synchronous progression to the S phase after HU treatment similar to results observed by Cools *et al* (2010). However, HC has a delayed and cumulative effect on cell proliferation from 16h post-treatment. This study did not observe the effect of HC beyond 24h. Therefore, following the effect of HC on root meristem cells for a longer duration will help get a better understanding of how HC affects the cell cycle. Overall, HC affects cell proliferation in a different manner relative to HU by blocking G2-M transition and inhibiting mitosis as observed earlier in onion roots treated with slightly higher concentrations of HC which caused increased oxidative stress seen as accumulation of ROS (Soltys *et al*., 2011).

Intracellular redox is highly regulated at major cell cycle checkpoints, G1 and G2, to ensure proper progression of cells through the cell cycle (De Simone *et al*., 2017). Inter-compartmental transport and sequestration of the antioxidant glutathione is pivotal in modulating the cellular redox state (De Simone *et al*., 2017). Treatment with HC in this study triggered an oxidation event in the cytosolic compartment immediately post-treatment causing an increase in redox potential similar to the effect observed on HU treatment (De Simone, 2016) due to the transport of GSH into the nucleus from the cytosol during G1 phase leading to depletion of the cytosolic GSH pool indicated by the higher degree of oxidation in the cytosol compared to the nuclei (Diaz-Vivancos *et al*., 2010b). The HU treated synchronized cells were predominantly in G1 phase at this time period (Cools *et al*., 2010; De Simone, 2016). However, the reason behind the similar reaction by HC treated unsynchronized cells cannot be clearly understood. HC treatment did not cause any significant alteration in the redox potential of the nucleus in the early hours of HC treatment but at the later hours of the treatment, when G2/M phase cells began to accumulate, both the nucleus and the cytosol of the cells were highly oxidised compared to the control and in stark contrast to HU treated cells (predominantly in G2/M phase), which were maintained in a reduced state (Cools *et al*., 2010; De Simone, 2016). The accumulation of cells at G2/M checkpoint at the later points of HC treatment could be due to the increased oxidation which depleted the cellular GSH pool causing GSH deficiency which alters the levels of CYCs and CDKs necessary for G2-M transition as observed in cucumber roots treated with 0.25 mM phenylcarboxylic acid which caused inhibition of *CYCB* gene expression (Inzé and De Veylder 2006; Gutierrez, 2009; Zhang *et al*. 2009). However, the reason for the delay in replenishing the GSH pool to overcome the inhibition in the absence of phytotoxicity is not clearly understood. This needs to be explored further. The cell cycle does not seem to be synchronized by HC in the 24h treatment period used in this study.

## Conclusion

The nuclei and cytosol of proliferation zone cells in Arabidopsis radicles are maintained in a highly reduced state and have similar glutathione redox potentials. HC treatment triggered increased oxidative stress towards the final hours of treatment which was accompanied by gradual accumulation of G2/M and depletion of G1/S phase cells possibly due to G2/M phase cell cycle arrest. This arrest could be due the depletion of total cellular GSH pool causing significant oxidation in both the nuclei and the cytosol. HC at a concentration of 1.5mM did not synchronize the cell cycle of root meristematic cells like HU in the 24h treatment period.

## Methodology

Unless otherwise stated, all chemicals were sourced from Sigma Aldrich.

### Plant material

*Arabidopsis thaliana* ([L.] Heynh.) wild type (Col-0) seeds expressing redox sensitive green fluorescent protein (roGFP2; Meyer *et al*., 2007) used to determine the redox state of the nuclei and cytosol in the embryonic root proliferation zone and wild-type (Col-0) seeds, were provided by Prof Christine Foyer (University of Leeds, UK) and Col-0 seeds of dual-core marker system (cell cycle tracking in plant cells; Cytrap) expressing *pCYCB1∷CYCB1-GFP and pHTR2∷CDT1a (C3)-RFP* (Yin *et al*., 2014) used to simultaneously monitor S/G2 and G2/M phases of cell cycle, were obtained from Dr Masaaki Umeda (Nara Institute of Science and Technology, Japan).

### Growth conditions

Col-0 (wild type), roGFP2 and Cytrap seeds were surface-sterilized and transferred to plates containing half-strength Murashige and Skoog agar medium (½MS; Murashige and Skoog, 1962), prepared from 2.2gL^−1^ MS basal medium, 0.5gL^−1^ 4-Morpholineethanesulfonic acid, 0.1gL^−1^ Myoinositol, 10gL^−1^ sucrose and 10gL^−1^ agar at pH 5.7. The seeds were then stratified at 4°C for 48h and allowed to germinate at 21°C in the dark for a further 48h. The 2d old seedlings were transferred to fresh ½MS medium in the absence (control) or presence (chemical treatments) of 3mM hydroxyurea (NH_2_CONHOH; HU, De Simone *et al*., 2017) or 1.5mM HC (H_2_CN_2_) and grown at 21°C in the dark until analysis (**Supplementary Figure S1**).

### Root growth rate

Root length of Col-0 (wild-type) seeds was measured 0, 24 and 48h after treatment with 3mM HU and 1.5mM HC and in control seeds using ImageJ image analysis software (Schneider *et al*., 2012) and the root growth rate was calculated. 3 biological replicates of 10 seeds each was used per time point for all the treatments.

### Visualization of redox status

The proliferation zone cells were identified based on the observations made by De Simone (2016) using PLT3∷GFP, WOX5∷GFP and WOL∷GFP markers to identify columella, QC and vascular system cells in embryonic roots of Arabidopsis at the same developmental stage as used in this study (Figure 1, 2).

**Figure 1.**
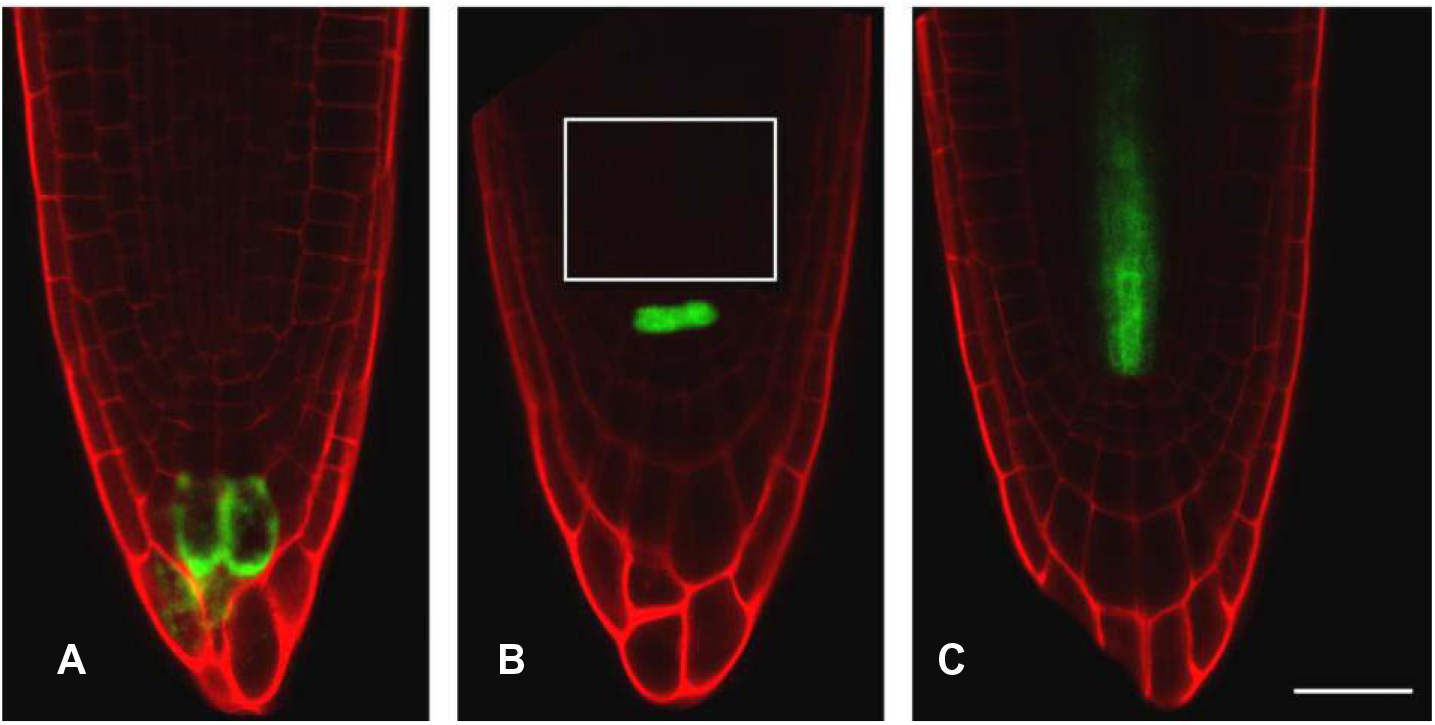
Distribution of GFP-tagged markers in the different cell types in *Arabidopsis thaliana* roots. Arabidopsis root tips expressing GFP tagged to (A) PLETHORA (PLT) gene encoding AP2-domain transcription factors (PLT3), (B) WUSCHEL-related homeobox 5 (WOX5) (white box marks the proliferation zone) and (C) WOODEN LEG (WOL). Roots were stained with PI on a microscope slide. Scale bar = 25 μm. (Reproduced from De Simone, 2016).

**Figure 2.**
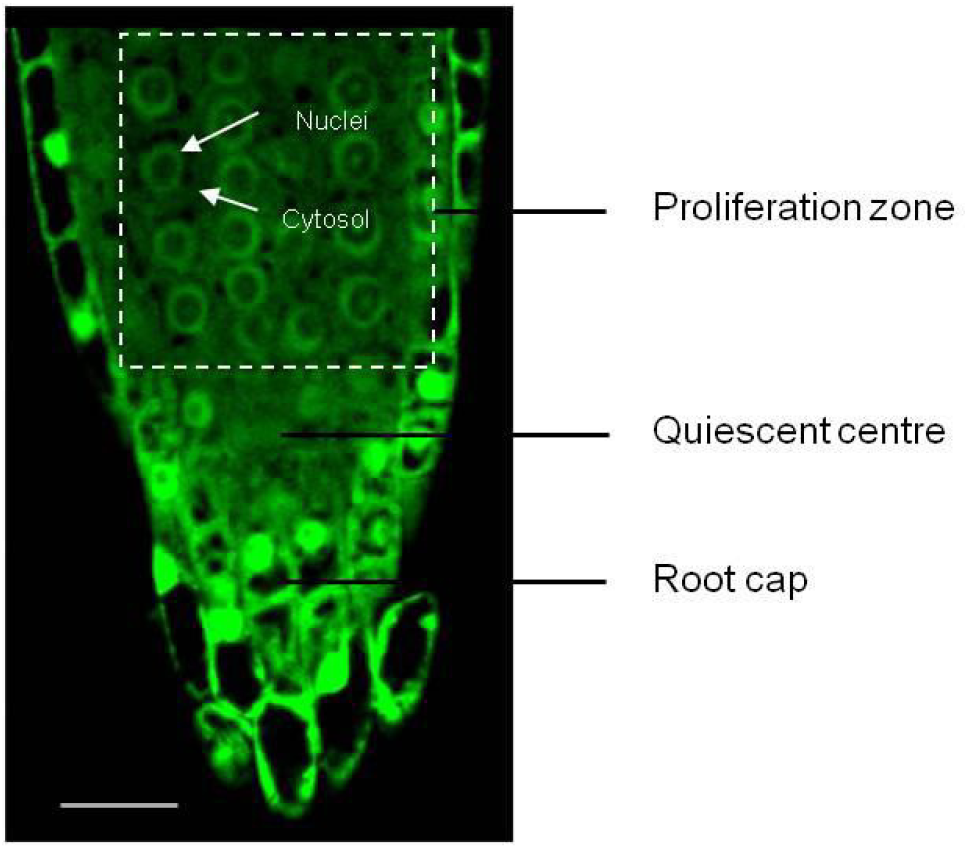
Root zones in *Arabidopsis thaliana* embryonic root imaged using confocal microscope at excitation wavelength of 488nm. Proliferation zone, quiescent centre and root cap cells in an Arabidopsis embryonic root identified based on observations made by De Simone (2016) (Figure 1). The white arrows indicate nuclei and cytosol of a proliferation zone cell. Bar= 25 μm.

Redox measurements were carried out every alternate hour (i.e. 0h, 2h, 4h…) over a period of 24h after transfer to HC and control treatments as described in the previous section. Germinated roGFP2 seeds collected at various time points after treatment were placed on a drop of sterile water on a clean slide and imaged using the 40X/1.3 Oil DIC M27 lens of LSM700 Carl Zeiss inverted confocal microscope (Carl Zeiss AG, Germany) at excitation wavelengths of 405nm for the oxidized and 488nm for the reduced form of roGFP2 (De Simone *et al*., 2017; Figure 3). The degree of oxidation in the nuclei and cytosol of the cells in the proliferation zone above the quiescent centre was later determined from the ratio of the fluorescence intensities at 405 and 488nm (405nm:488nm) measured using ImageJ image analysis software (http://rsbweb.nih.gov/ij/). 5 technical and 5 biological replicates per time point were used in this study.

**Figure 3.**
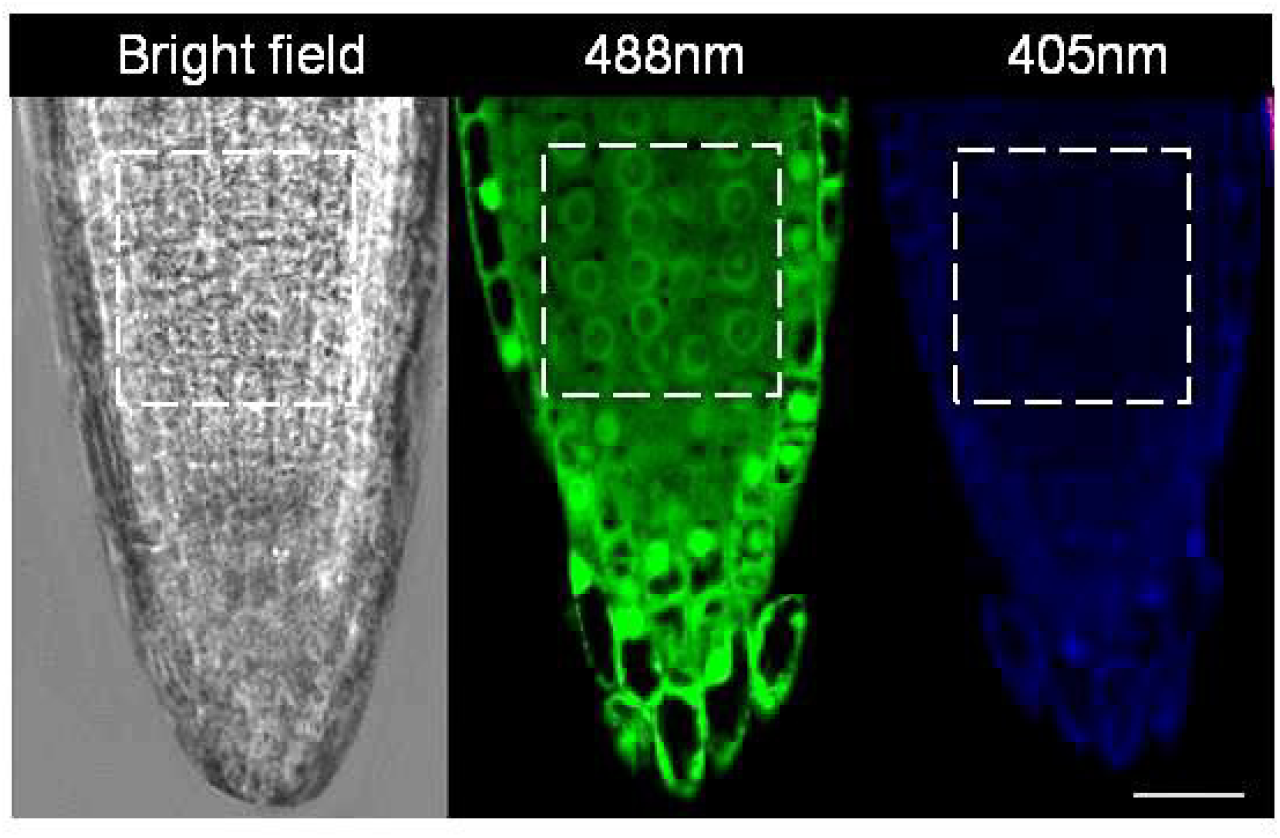
Confocal images of *Arabidopsis thaliana* embryonic root tip in 2d old roGFP2 seedlings. The boxed area indicates the proliferation zone where fluorescence intensity measurements were carried out. Bar = 25μm.

### Calibration of roGFP2 probe

The roGFP2 probes were calibrated at the end of the experiment as described by De Simone *et al*. (2017). The embryonic roots were immersed in 2mM dithiotheritol (DTT) solution for 10 min to cause complete reduction of roGFP2 before being imaged at excitation wavelengths of 405nm and 488nm. Similarly, to completely oxidize the roGFP2 probe, the embryonic roots were treated with 2mM hydrogen peroxide (H_2_O_2_) solution for 15 min and later imaged at 405nm and 488nm as shown in Figure 3. The 405/488nm ratios calculated from these treatments were used for calibration of the roGFP probe during the calculation of the degree of oxidation and redox potential in the nuclei and cytosol of the cells in the proliferation zone of the embryonic root tip as described in the next section.

### Calculation of redox potential

Oxidation degree (**OxDroGFP2**) and redox potential of roGFP2 (**EroGFP2**) probe were calculated as described by Meyer *et al*. (2007)

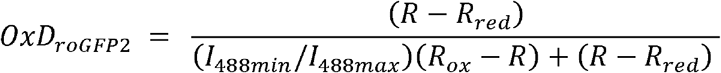

**OxDroGFP2** was obtained using the roGFP2 fluorescence ratio (405/488 nm ratio) obtained using confocal microscopy as described previously, where, **R** is the ratio of excitation at 405 and 488nm, **Rred** is the ratio of fully reduced roGFP2 obtained from 2mM DTT treated embryonic root tip cells, **Rox** is the ratio of fully oxidized roGFP2 obtained from 2mM H_2_O_2_ treated embryonic root tip cells and **R** is the fluorescence ratio (405/488nm ratio). **I488min** and **I488max** are the fluorescence intensities of completely reduced and completely oxidized roGFP2 measured with excitation at 488nm (Figure 3).

**OxD roGFP2** obtained from the above equation was used to estimate the glutathione redox potential (**EGSH)** in mV using the following Nernst equation,

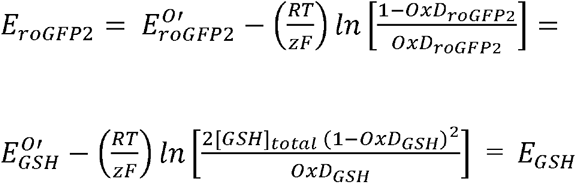

where, **E** is the redox potential and **EO**’ is the midpoint potential of GFP and roGFP2, **R** is the gas constant (8.315 JK^−1^mol^−1^), **T** is the absolute temperature (298.15 K), **z** is the number of electrons exchanged (2) and **F** is the Faraday constant (9.648 × 10^4^ Cmol^−1^).

### Cell cycle status

Cell cycle was monitored using Cytrap system (*Yin et al., 2014)* and flow cytometry.

#### Flow cytometry

Intact nuclei suspension was prepared from fresh embryonic root tips using slight modifications to the protocol used by Arumuganthan and Earle (1991). ~100 seeds per biological replicate (3 biological replicates) were used. Embryonic root tips of control and HC treated Col-0 wild type seeds were carefully collected at 0, 5, 12, 16 and 24h after treatment and chopped with a razor blade in ice cold nucleus-isolation buffer at pH 7.4 (10mM MgSO4·7H2O, 50mM KCl, 5mM HEPES, 1mg.mL^−1^ DTT, 0.5%v/v TritonX-100 and 1% (w/v) PVP-40) on ice and incubated on ice for 1h with gentle swirling every 1/2h. Following incubation, the suspension was passed through a 40μm nylon mesh, centrifuged at 100*xg* for 10min at 4°C and the supernatant was carefully discarded. Later, the nuclei pellet was resuspended in 2mL nucleus-isolation buffer and treated with RNase for 6min at room temperature before being stained with 20μg.mL^−1^ propidium iodide and stored on ice until analysis. Stained cell sample was then run on BD FACSCalibur flow cytometer (BD biosciences, Europe) at the Centre for Microscopy Characterisation and Analysis; CMCA, UWA, equipped with a primary blue 488nm laser and data for ~100,000 nuclei were recorded (i.e. until 20,000 G2 (4C) nuclei were collected). The proportion of nuclei with 2C and 4C DNA content was recorded. The data were visualised real-time using scatter dot plots (FSC and SSC) and histograms. The results were analysed using the Flowing Software version 2.5 (http://flowingsoftware.btk.fi/) by manual gating to eliminate debris from the population of interest on the scatter plots and subsequent generation of histograms from the scatter plot data in which the different populations were gated to obtain the final proportion of G1, S and G2 nuclei computed by the software. Later the data was plotted using Microsoft Excel 2016.

#### Cytrap

The Cytrap dual cell cycle phase marker system was used to monitor both S/G2 and G2/M phases of the cell cycle simultaneously in proliferating cells of embryonic root (Yin *et al*., 2014). The root tips of control and HC treated Cytrap seeds placed in a drop of water on a clean slide were imaged at 40X magnification using the LSM700 Zeiss inverted confocal microscope at excitation wavelengths of 488nm for GFP (Figure 4A) and 559nm for the RFP (Figure 4B) at 0, 2, 5, 8, 10, 16, 18 and 24h after treatment.

**Figure 4.**
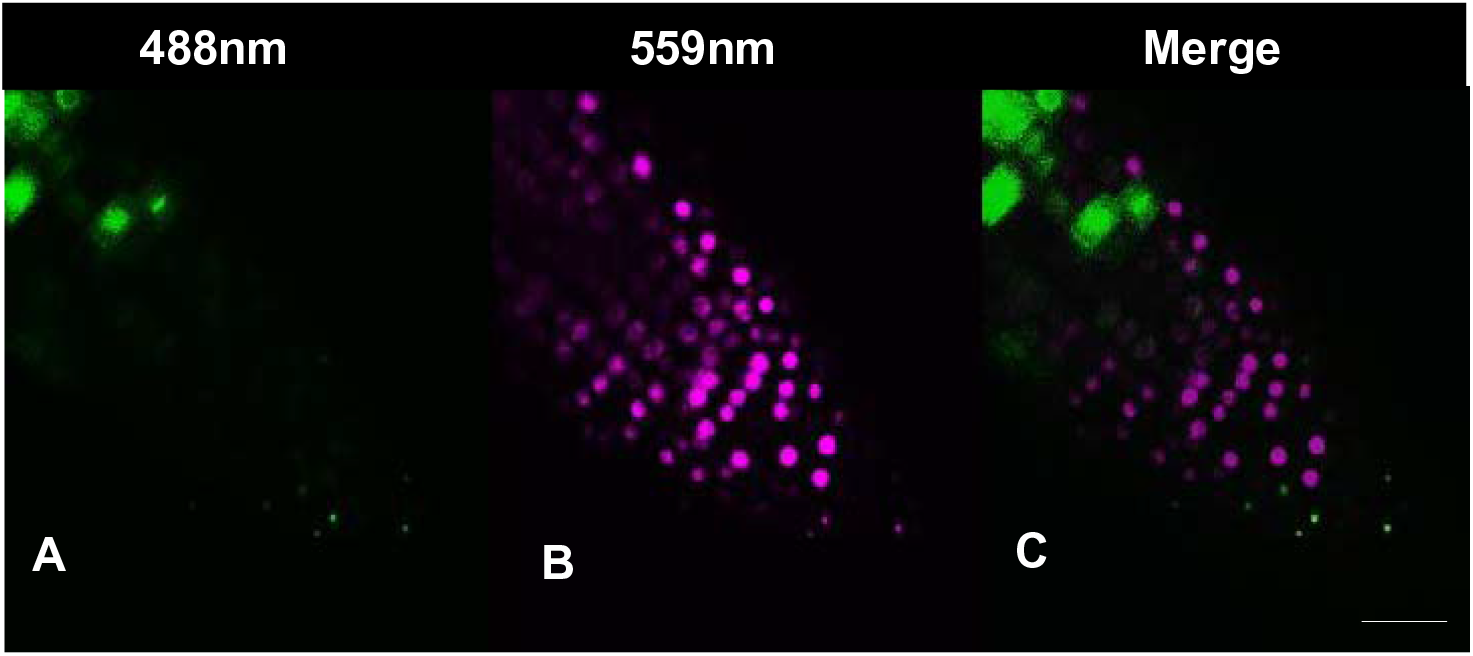
Confocal images of *Arabidopsis thaliana* pHRT2∷CDTa (C3)-RFP (magenta) and pCYCB1∷ CYCB1-GFP (green) root tip obtained from excitation at 488 (GFP) and 559nm (RFP). (A) G2/M phase cells (green); (B) S/G2 phase cells (C3)-RFP (magenta); (C) Merge of A and B. Bar=25μm.

**Figure 5.**
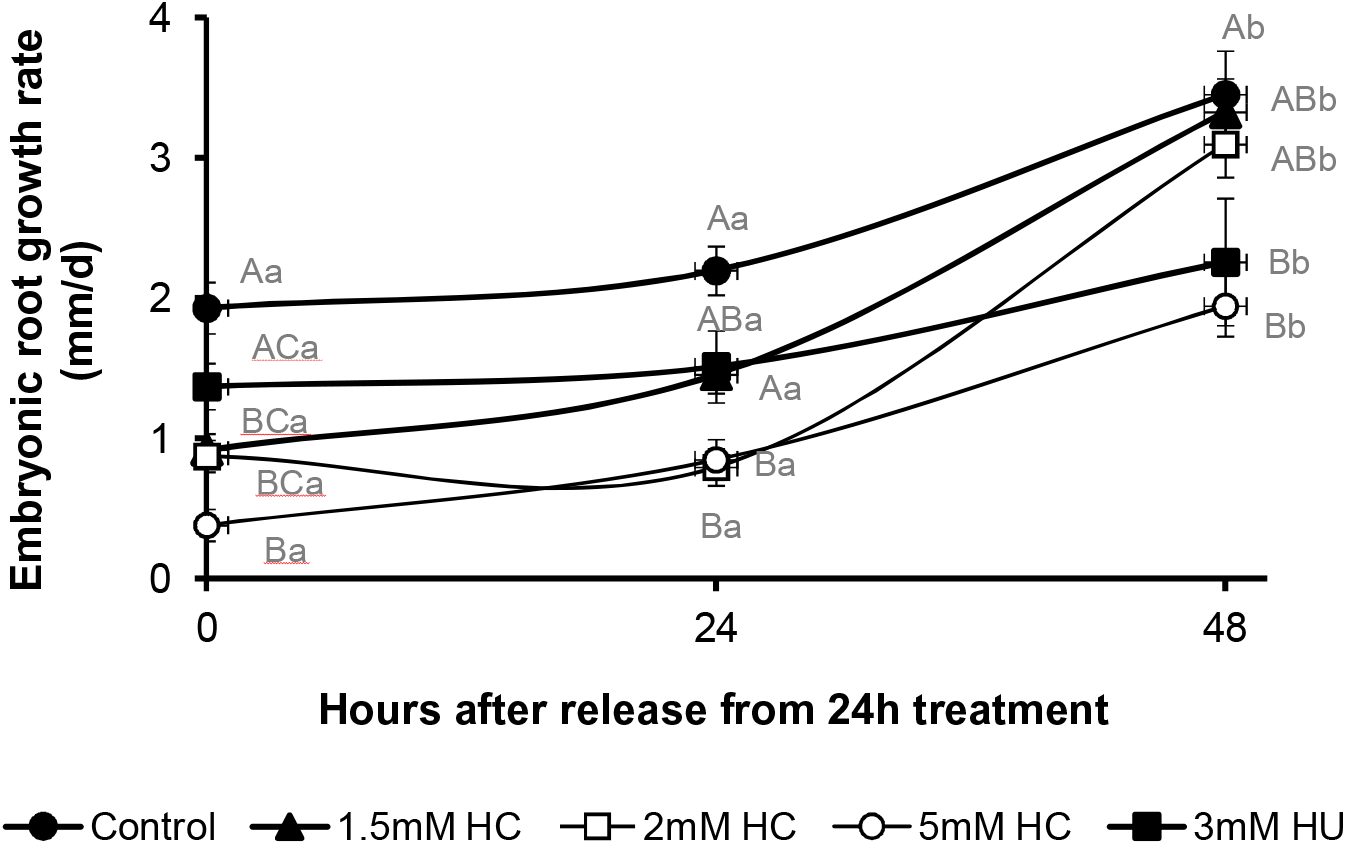
Recovery of root growth after release from treatment in control, HC and HU treated *Arabidopsis thaliana* seedlings. Rate of root growth in control, 1.5, 2 and 5mM HC and 3mM HU treated seedlings 0, 24 and 48h after recovery from 24h treatment in the dark at 21°C. Lower-case letters above bars denote significant differences (p◻≤0.01) within same treatment at different time points and upper-case letters above bars denote significant differences (p◻≤0.01) at same time point among different treatments corroborated using Tukey’s honestly significant difference (HSD) test.

**Figure 6.**
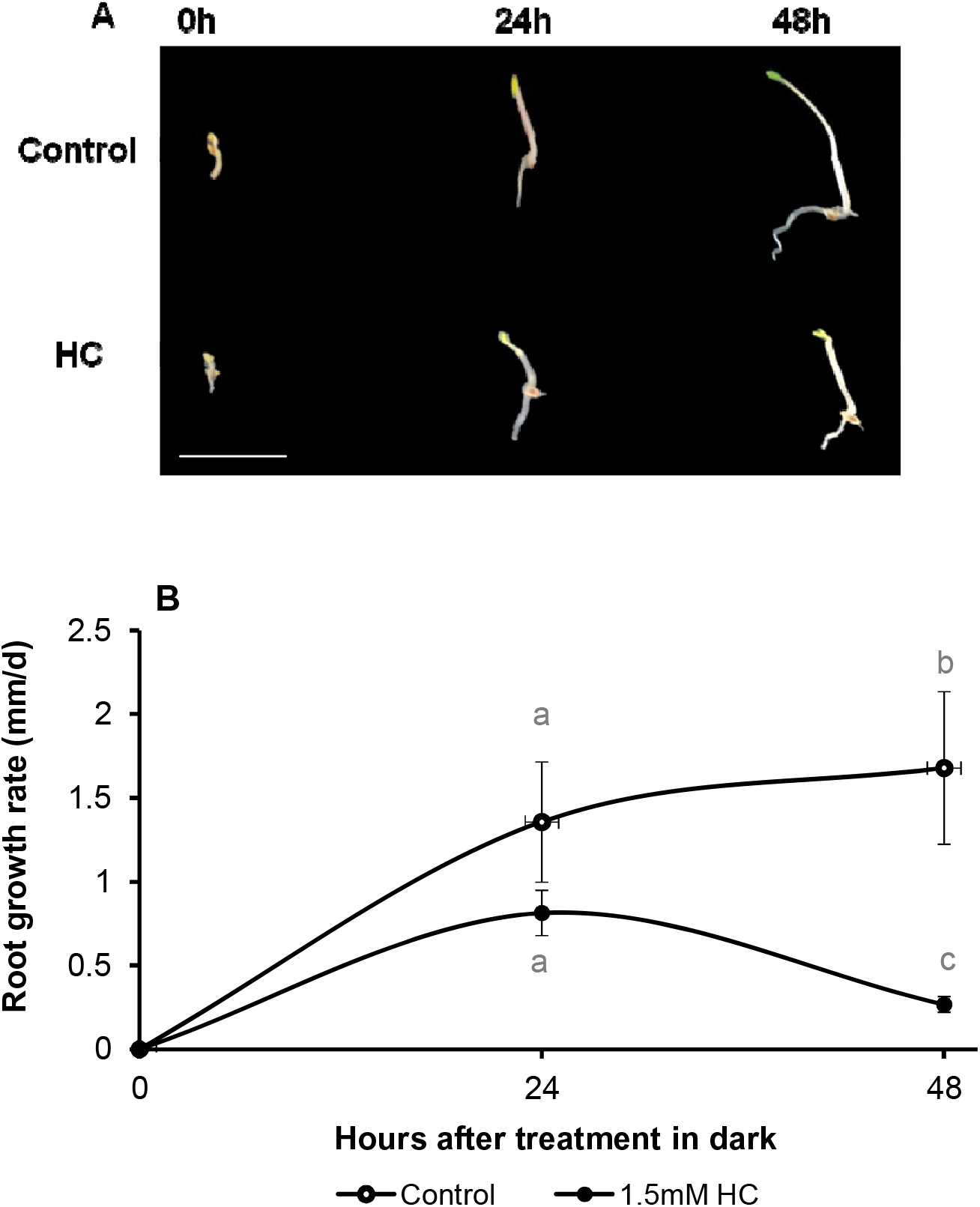
Root growth in control and HC treated *Arabidopsis thaliana* seedlings. (A) Macroscopic images of control and HC treated seedlings after 0h, 24h and 48h of treatment. Bar=1cm. (B) Rate of root growth in control (open circles), 1.5mM HC (closed circles) treated seedlings after 24 and 48h of treatment in the dark at 21°C. Lower-case letters above bars denote significant differences (p◻≤0.01) at same time point among different treatments corroborated using Tukey’s honestly significant difference (HSD) test. There was no significant difference at p◻≤0.01 within same treatment at different time points.

**Figure 7.**
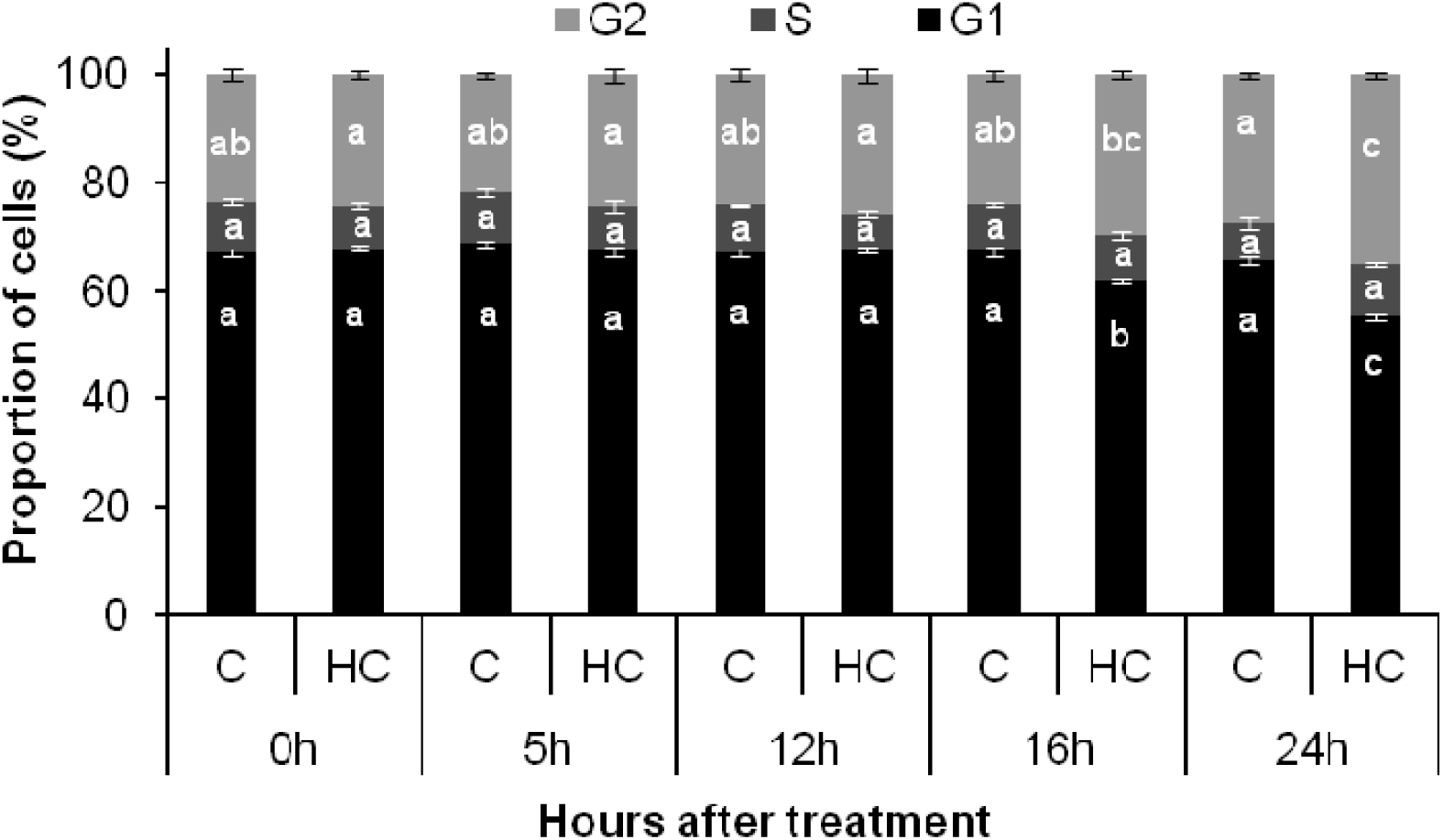
Distribution of cells in each cell cycle phase in control and HC treated root tips. Proportion of cells in G1, S and G2 phases of the cell cycle in control and HC treated Arabidopsis embryonic root tip cells at various time points of treatment. Lower case letters below the bars denote significant differences (p◻≤0.01) in the distribution of cells within the same cell cycle phase among different time points of control and HC treatment corroborated using Tukey’s honestly significant difference (HSD) test.

**Figure 8.**
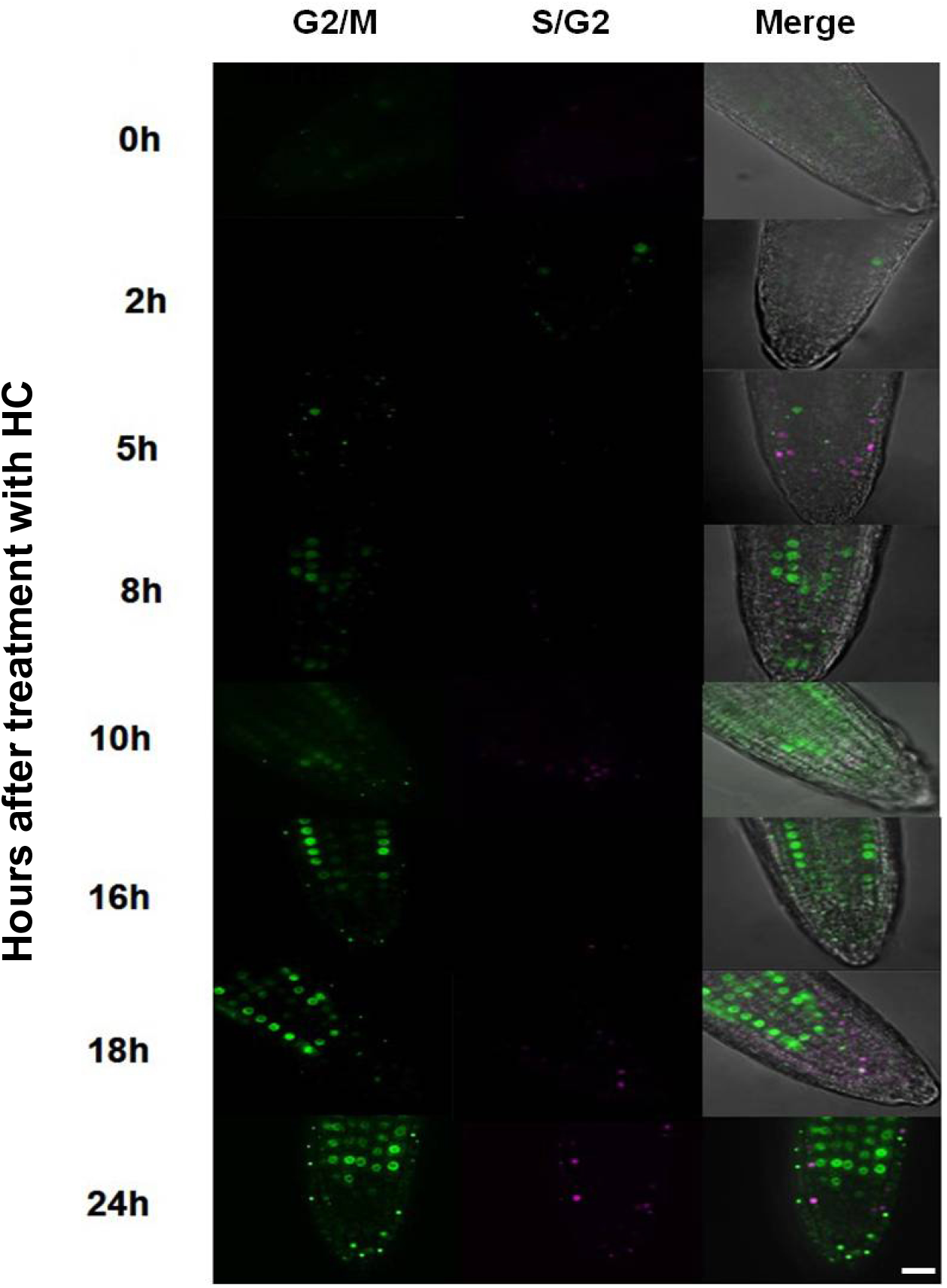
Cytrap expression in control and HC treated Arabidopsis embryonic root tip cells at various time points of treatment. The expression pattern in the control remained the same at all time points and similar to 0h treatment of HC. Hence, a common representative figure for the control and 0h HC treatment is shown here. Magenta and green fluorescence shows distribution of cells in S/G2 (Phtr2∷ CDT1a (C3)-RFP expression) and G2/M (Pcycb1∷ CYCB1-GFP expression) phase of the cell cycle respectively and merge shows overlay of S/G2 and G2/M with a bright-field image background. Bars= 25μm. Only one representative figure out of 3 replicates for each time point is shown.

**Figure 9.**
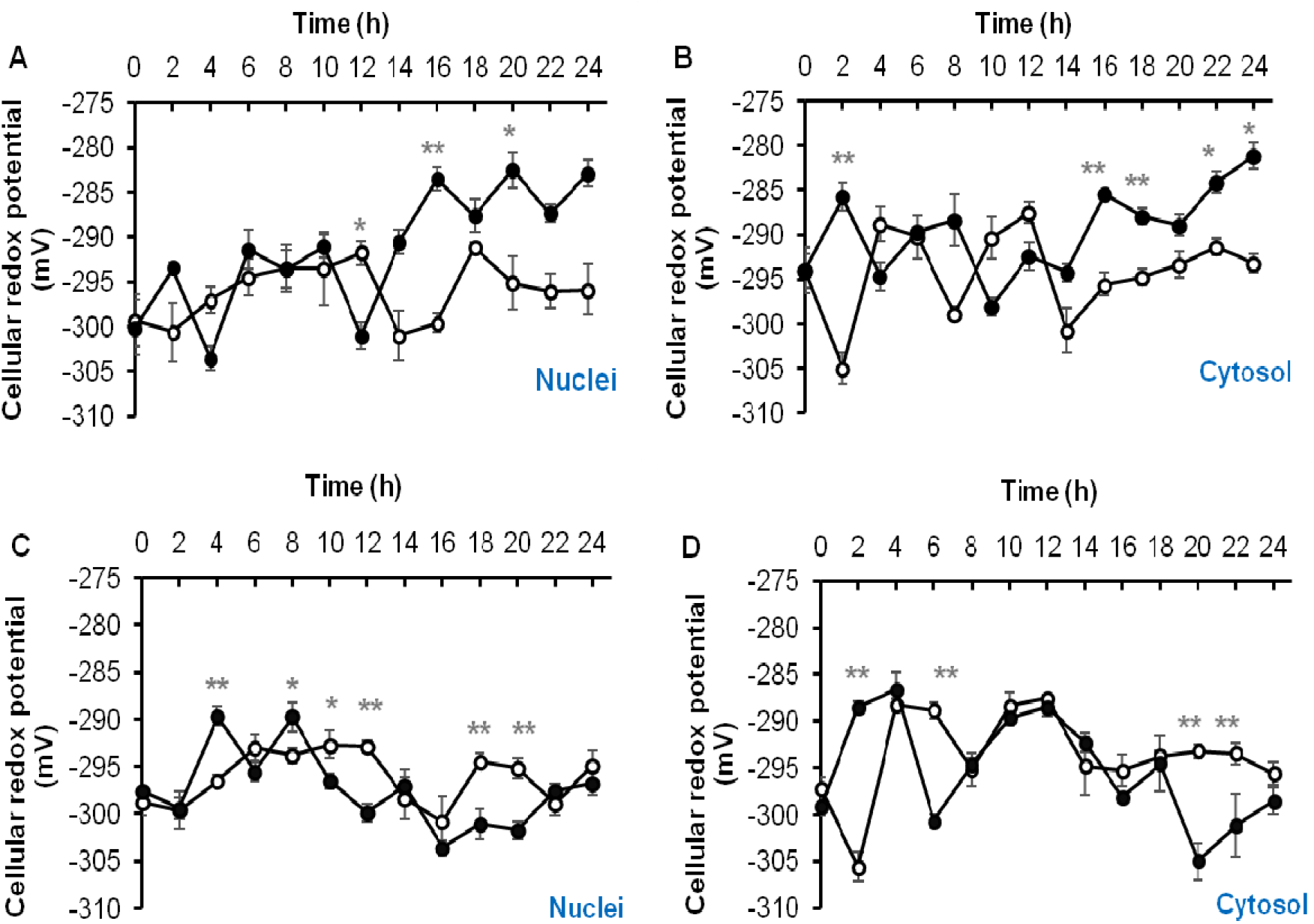
Changes in cellular redox potential occurring in the proliferating cells of embryonic root of germinating *Arabidopsis thaliana* seeds. Effect of HC (A, B) and HU (C, D) treatment on cellular redox potential of meristematic zone cells, in roGFP2-expressing roots, during 24h of treatment. Fluorescence was measured in the absence (open circles) or presence (filled circles) of treatment with HC and HU. * and ** above bars denote significant differences (*p◻<◻0.05, **p◻<◻0.01) in comparison to the untreated control, using Tukey’s honestly significant difference (HSD) test.

**Figure 10.**
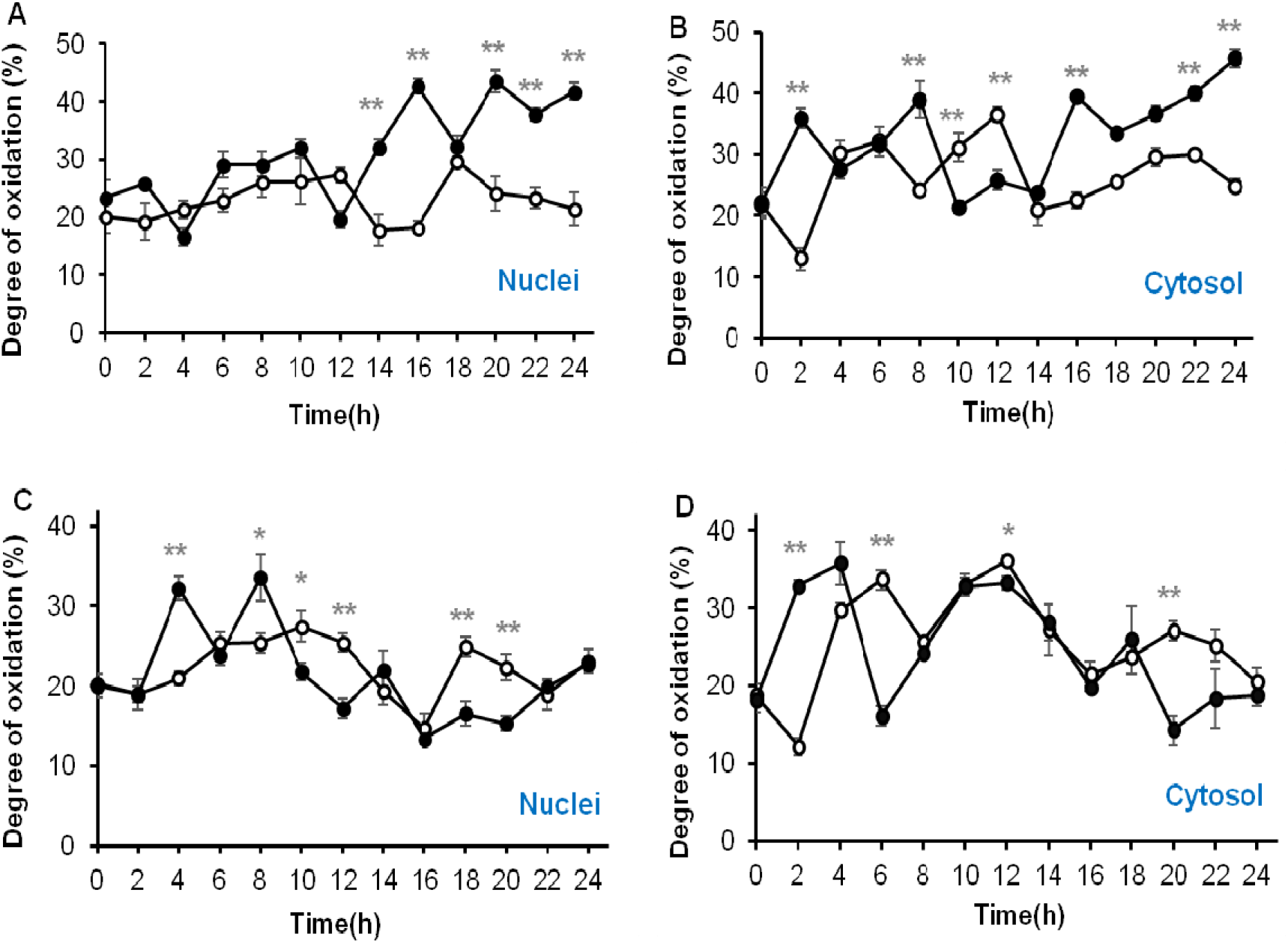
Changes in cellular oxidation occurring in the proliferating cells of embryonic root of germinating *Arabidopsis thaliana* seeds. Effect of HC (A, B) and HU (C, D) treatment on degree of oxidation of meristematic zone cells, in roGFP2-expressing roots, during 24h of treatment. Fluorescence was measured in the absence (open circles) or presence (filled circles) of treatment with HC and HU. * and ** above bars denote significant differences (*p◻<◻0.05, **p◻<◻0.01) in comparison to the untreated control, corroborated using Tukey’s honestly significant difference (HSD) test.

### Data analysis and statistics

All calculations were performed and graphics were compiled using Microsoft Excel 2016. Significant differences among various sampling dates were corroborated statistically by applying one-way ANOVA test, using Tukey’s honestly significant difference (HSD) posthoc test with P≤0.01 and P≤0.05 (Origin; OriginLab, Northampton, MA).

## Acknowledgements

We are very grateful to the team of the Centre for Microscopy, Characterisation and Analysis of The University of Western Australia for their technical guidance. We also acknowledge support and teamwork of other laboratory members.

## Author contributions

YV performed all the experiments, analysed the data, prepared all the figures, and drafted the manuscript with constructive comments from co-authors. CF and ADS supervised roGFP experiments, ADS performed Fig. 1. MJC and CF conceived and supervised the project. SS and JAC assisted calibration and interpretation. All authors contributed to the article and approved the submitted version.

## Data availability

Data are available from the corresponding author, Michael Considine, upon request.

---

**Supplementary Figure S1. Schematic showing the experimental design**. roGFP2 and Cytrap seeds were stratified for 48hr in the dark at 4°C and later allowed to germinate at 21°C under dark condition for 48hr, followed by transfer to chemical treatment (3mM HU or 1.5mM HC) or control conditions for root growth measurements, *in vivo* measurement of redox state and monitoring cell cycle status in the proliferation zone of the embryonic root. HC-Hydrogen cyanamide, HU-Hydroxy urea.

**Supplementary Figure S2. Cytrap expression in control Arabidopsis embryonic root tip cells at various time points of treatment**. The expression pattern in the control remained the same at all time points. Magenta and green fluorescence shows distribution of cells in S/G2 (Phtr2∷CDT1a (C3)-RFP expression) and G2/M (Pcycb1∷ CYCB1-GFP expression) phase of the cell cycle respectively and merge shows overlay of S/G2 and G2/M with a bright-field image background. Bars= 25μm. Only one representative figure out of 3 replicates for each time point is shown.

