## Supplementary fig file for "Hydrogen Cyanamide Causes Reversible G2/M Cell Cycle Arrest Accompanied by Oxidation of the Nucleus and Cytosol in *Arabidopsis thaliana* Root Apical Meristem Cells"

Supplementary Figures

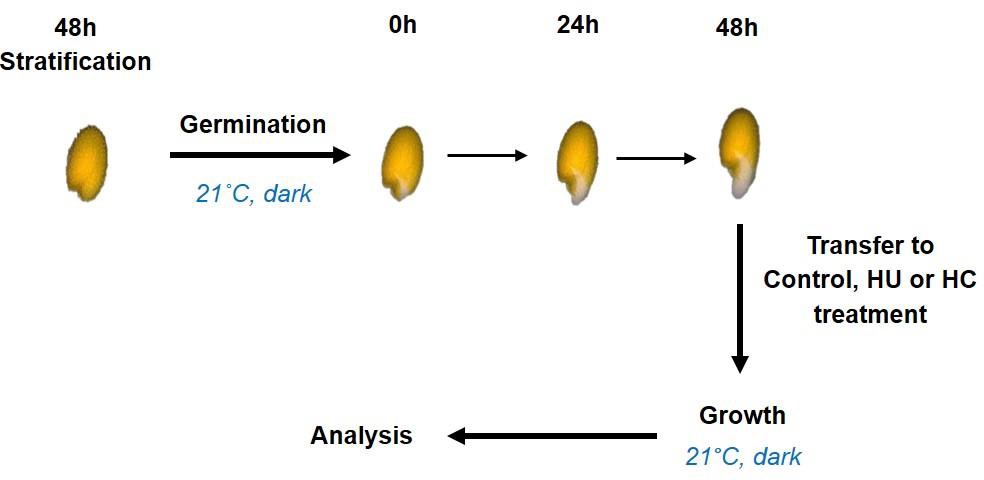

**Supplementary Figure S1. Schematic showing the experimental design.** roGFP2 and Cytrap seeds were stratified for 48hr in the dark at 4˚C and later allowed to germinate at 21˚C under dark condition for 48hr, followed by transfer to chemical treatment (3mM HU or 1.5mM HC) or control conditions for root growth measurements, *in vivo* measurement of redox state and monitoring cell cycle status in the proliferation zone of the embryonic root. HC- Hydrogen cyanamide, HU- Hydroxy urea.

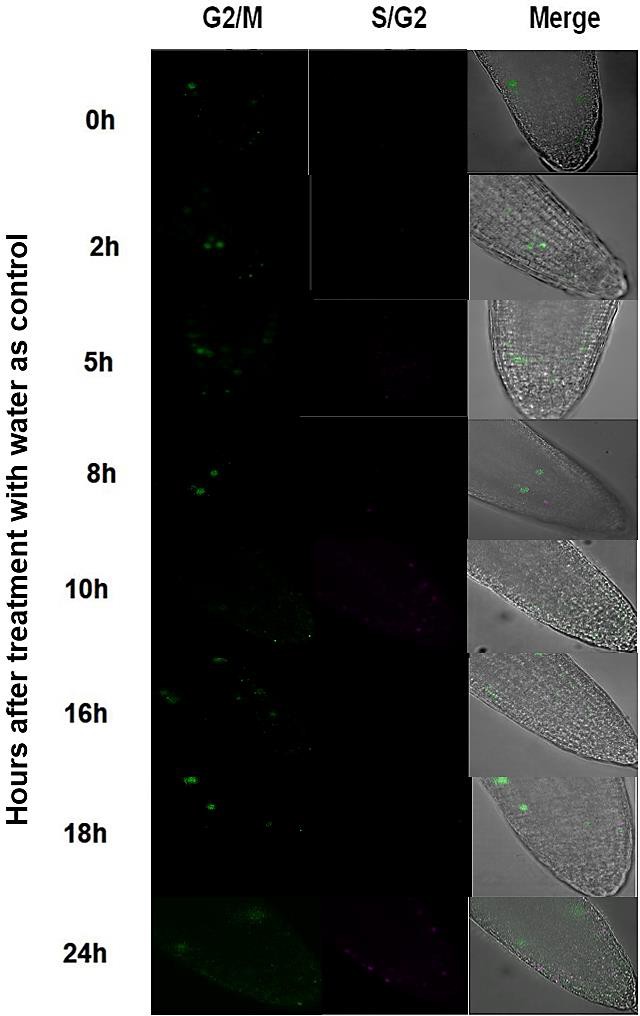

**Supplementary Figure S2. Cytrap expression in control Arabidopsis embryonic root tip cells at various time points of treatment.** The expression pattern in the control remained the same at all time points. Magenta and green fluorescence shows distribution of cells in S/G2 (Phtr2::CDT1a (C3)-RFP expression) and G2/M (Pcycb1:: CYCB1-GFP expression) phase of the cell cycle respectively and merge shows overlay of S/G2 and G2/M with a bright-field image background. Bars= 25µm. Only one representative figure out of 3 replicates for each time point is shown.
